# Analysis of glue line and correlations between anatomical characteristics of *Eucalyptus grandis* × *Eucalyptus urophylla* glued-laminated timber

**DOI:** 10.1101/599662

**Authors:** Rafael G. E. Oliveira, Fabricio G. Gonçalves, Pedro G. A. Segundinho, José Tarcísio S. Oliveira, Juarez B. Paes, Izabella Luzia S. Chaves, Alice S. Brito

## Abstract

The main goal of this study was to analyze glue line on eucalyptus wood. In order to do that, thickness of main and secondary glue lines were measured as well their interaction with apparent density of elements glued with resorcinol-formaldehyde (RF) and castor polyurethane (CP) adhesives. Anatomical wood characterization of *Eucalyptus grandis* × *Eucalyptus urophylla* was performed by correlating glue line thickness. According to normative instruction, specimens were produced for delamination tests. The experiment was conducted in a completely random 2 × 2 design factorial scheme (two classes of apparent density and two adhesives). Pearson correlation (t < 0.01) was performed among variables. It was found that there was adhesive penetration into wood pots and rays. Glue line thickness was higher in woods with density higher than 0.58 g cm^−3^ glued with RF adhesive. There was low correlation among density, vessel diameter, main and secondary glue lines (t < 0.01).

## Introduction

Glued laminated timber (Glulam) is a structural product obtained by gluing pieces of timber with fibers parallel to each other [1]. The market still needs to know more information about the mechanical resistance of this product; for that reason, to understand how glulam will work it is important to evaluate the behavior of some variables, such as, apparent density and timber glued with adhesive.

Wood density provides information to support methods that should be adopted during the gluing process. By its determination, it is possible to correlate adhesion resistance with anatomic elements. Proportion of empty spaces combined with dimensions and arrangement of cellular elements have influence on adhesive’s mobility and penetration into timber structure as well in the resistance of glue line to delamination [2, 3, 4, 5, 6, 34]. When low density (higher frequency and vessel diameters, with high and wide radii), it may allow excessive penetration of adhesive, if it does not present an ideal viscosity and formation of ravenous glue line [7, 8].

When species are anatomically unfavorable for gluing, there will be low adhesive penetration and formation of a thick glue line [8, 9]. Both types of glue lines are undesirable, because they reduce mechanical strength of glued joints causing separation of adjacent layers. Variations on adhesive viscosity can correct this problem [3].

Physico-chemical adhesion phenomenon predicts an interaction mechanism between solid surfaces glued by adhesive and the capacity of holding other materials together [10]. Adhesives are used to join elements by flowing and filling empty spaces between them. Thus, they can reduce distances and create interactions among glued elements [11].

It is important to understand the interaction between wood and adhesive, because it helps to evaluate glueing quality [7], Especially when it comes to wood of *Eucalyptus* genus, since its anatomical structure presents occurrence of small diameter, presence of tilose, low frequency and small vessel width. Other characteristics are also important, such as the frequency of conducting vessels (anatomical parameter that most influences gluing process), followed by frequency, width of rays and conducting vessels diameter [12].

Therefore, the objective of this study was to measure, thickness of main and secondary glue lines and to correlate this variable with apparent density, vessels and rays of *Eucalyptus grandis* × *Eucalyptus urophylla* wood.

## Materials and methods

### Origin and initial preparation of the material

The board shaped timber used came from 11-years-old *Eucalyptus grandis* × *Eucalyptus urophylla* clones from Bahia Wood Products, located in the city of Nova Viçosa, Bahia, Brazil. Fig. 1 show the preparation process of the glued laminated timber and the other tests.

**Fig. 1.**
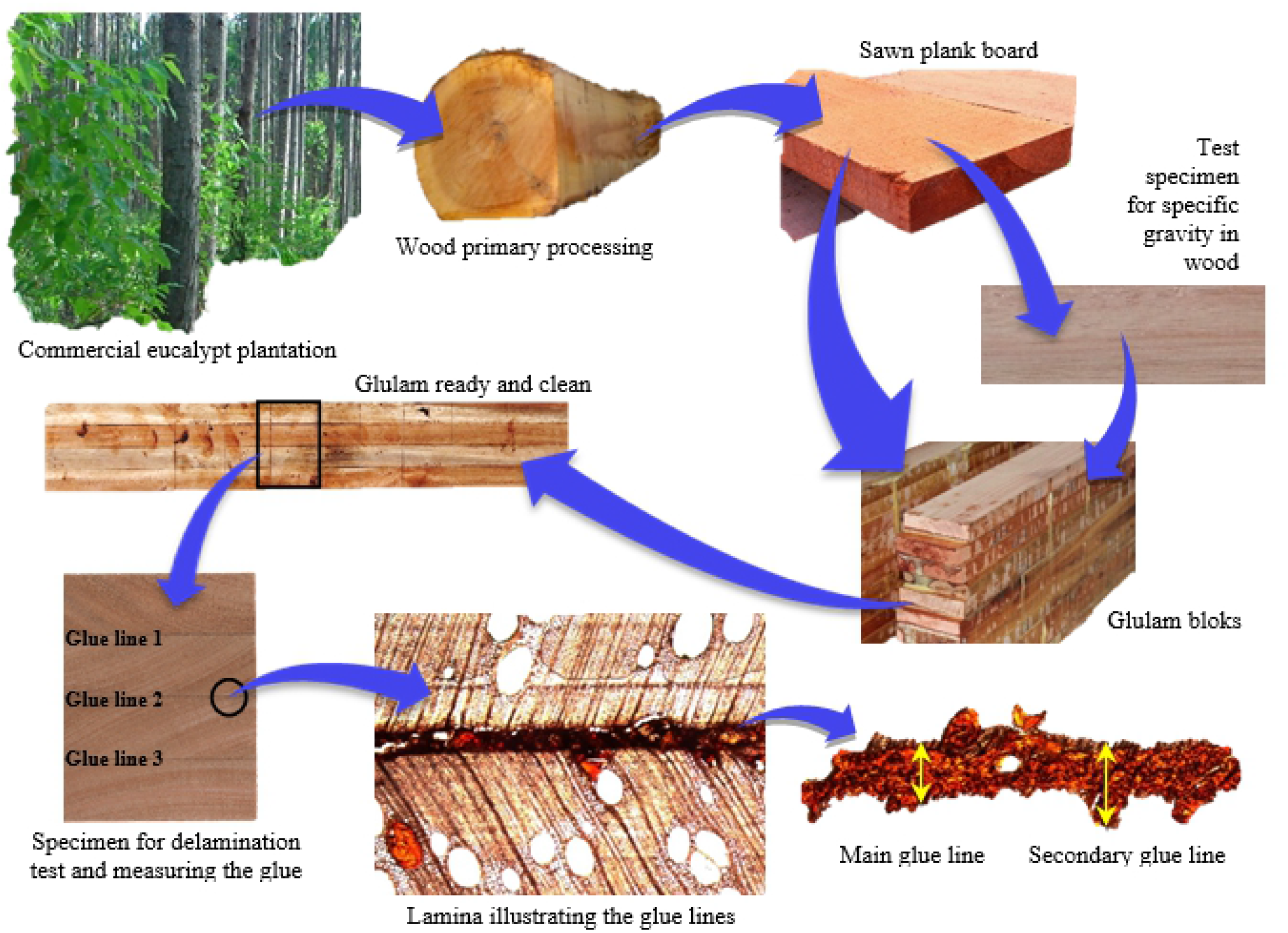
Method of obtaining timber boards and specimens.

There were 40 timber boards provided by the company, which were placed in a weather-protected environment until reaching the equilibrium humidity of the site. After that, the timber boards were visually classified and separated by desired dimensions to produce Glulam blocks.

The characterization of the wood physical properties (moisture content and apparent density) was performed according to Brazilian normative regulation NBR 7190 of the Brazilian Association of Technical Standards [13], using the same specimens.

To perform the tests, 4 specimens were removed from each timber board, totaling 160 specimens, which were sectioned in dimensions of 3 × 2 × 5 cm^3^ in the radial, tangential and longitudinal directions, respectively.

### Adhesive and glued elements

There were tested two commercial adhesives, a thermoset, the Cascophen RS-216-M based on resorcinol (RF), and a thermoplastic, the bi-component polyurethane based on castor oil.

Instructions of how to use the adhesives, as well the extender and catalyst choice were made based on recommendations from the manufacturer. For Cascophen RS-216-M application, it was added 20% of FM-60-M catalyst. The proportion 1:1.2 was used in the application of polyurethane, 1 part of component A (prepolymer) and 1.2 part of component B (polyol).

To produce glulam, timber boards were divided according to their apparent density into two groups: G1) apparent density less than 0.58 g cm^−3^; and G2) apparent density equal or higher than 0.58 g cm^−3^. After dividing the groups, timber boards were sectioned and transformed into 56 lamellae with dimensions of 21 × 21 × 2.5 cm^3^.

Each group had 28 lamellae, resulting in 14 Glulam elements. 7 Glulam blocks were used for each adhesive. A spatula was used to apply 300 g m^−2^ to each pair of timber board in a single glue line. They were joined and pressed in manual press (capacity 15 tons) for 48 hours at a pressure of 1 MPa and temperature of 20 °C [13].

### Wood Anatomical Characterization

The vessels (diameter and frequency) and rays (height, width and frequency) measurements were made from samples wood with 1.0 × 1.5 × 2.0 cm^3^ dimensions, in the radial, tangential and longitudinal directions, respectively, made by microtome.

Wood lamina were photomicrographed using the software AxioVision Rel. 4.5. In addition, it was performed microscopic identification using a 10x magnification lens [14].

### Visualization of the wood-adhesive interface

The study of wood-adhesive interface was carried out based on a methodology known [7]. The timber blades came from glued joints of a 0.5 × 0.5 × 0.5 cm^3^ dimension central hub, on a microtome. Each cube had samples collected in the glue line.

A total of 28 cubes were prepared, 14 cubes for each group of apparent density (7 cubes for each adhesive) resulting in 56 timber lamina. For each cube 30 measurements were taken, which means, 15 for the main glue line and 15 for the secondary glue line.

Thickness measurements were done in the main and secondary glue lines, through glue line width using a 5x objective and AxioVision Rel. 4.5 software.

### Delamination test on blocks of glued laminated timber

Delamination test was executed arranging the specimens inside an autoclave. After that, glue lines were exposed to stresses that came from vacuum and pressure effects. That is possible due to 6 days period of moistening and drying cycles - Steps 1 to 4 adapted from the American Institute of Timber Construction [15] (Fig. 2). Each cycle lasted 48 hours, out of a total three.

**Fig. 2.**
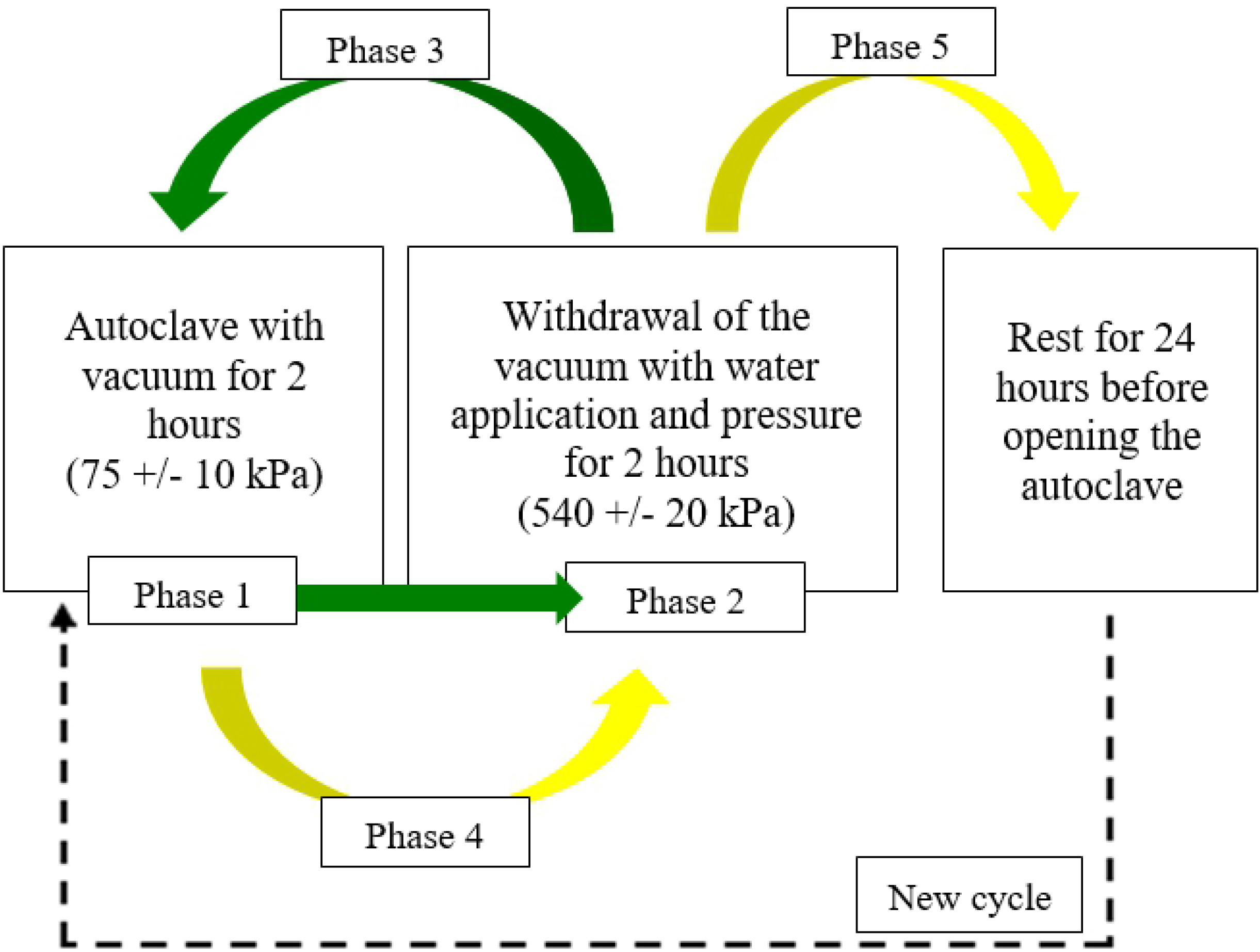
Scheme of moistening cycle for delamination test on glulam specimens. Adapted from the American Institute of Timber Construction [15].

At end of the third cycle, it was possible to tell, how Glulam would behave outside. This was done by measuring the delamination percentage gained in the two specimens’ top faces. It was possible to say the percentage of delamination, looking at the relation between maximum opening length and total length of the glue line.

Samples were placed outside in natural environmental conditions to dry out for 36 hours at 28 ± 2 ° C of temperature, aiming to reduce their weight, no more than 5 to 6% of the initial weight range. For each specimen, it is recommended that the total glue line delamination should not exceed 10% of total length at the specimen top, to glulam approval for exterior use [15].

### Statistical data analysis

Experiment was made in a completely randomized design following 2 × 2 factorial scheme, where apparent density and adhesive had two levels each one. Results were submitted to variance analysis (F < 0.05). For anatomical characteristics, it was also performed a descriptive quantitative statistical analysis.

Pearson correlation coefficient (t < 0.01) was used to find out the relationship between the following variables: apparent density, adhesive type, main glue line, secondary glue line, vessel diameter, ray height, width of rays, number of cells. Correlations were realized being 0.1 - 0.3 weak correlation, 0.4 - 0.6 moderate, 0.7 - 0.9 strong [16].

## Results

### Physical and anatomical wood characteristics

The hybrid *Eucalyptus grandis* × *Eucalyptus urophylla* showed an apparent average density (ρ) of 0.59 g cm^−3^ and an average moisture content of 9.34% (Table 1) These characteristics have influence on adhesive penetration [10], pressure need, pressing time and adhesive cure (how long it needs to be ready for use), making its determination an indispensable factor to produce glulam elements [32].

**Table 1.** Average and descriptive numbers of measured parameters in the wood rays of *Eucalyptus grandis* × *Eucalyptus urophylla*.

| Parameters evaluated | Density < 0.58 g cm<sup>-3</sup><br>(Group 1) | Density ≥ 0.58 g cm<sup>-3</sup><br>(Group 2) |
| --- | --- | --- |
| Height ($\mu\text{m}$) | 234.72 (63.91)<sup>a</sup> | 254.58 (73.69) |
| Width ($\mu\text{m}$) | 13.33 (2.93) | 16.18 (4.75) |
| Cells number in height | 14 (6) | 15 (3) |
| Frequency (rays mm<sup>-1</sup>) | 13 (2) | 16 (2) |

<sup>a</sup>Value in parentheses refers to standard deviation.

### Wood-adhesive interface and delamination test

Through photomicrographs, it was possible to see the adhesive penetration in the wood anatomical structure. A more visible glue line was found in the laminated timber glued with RF adhesive, due to its reddish color (Figure 3C and 3D).

**Fig. 3.**
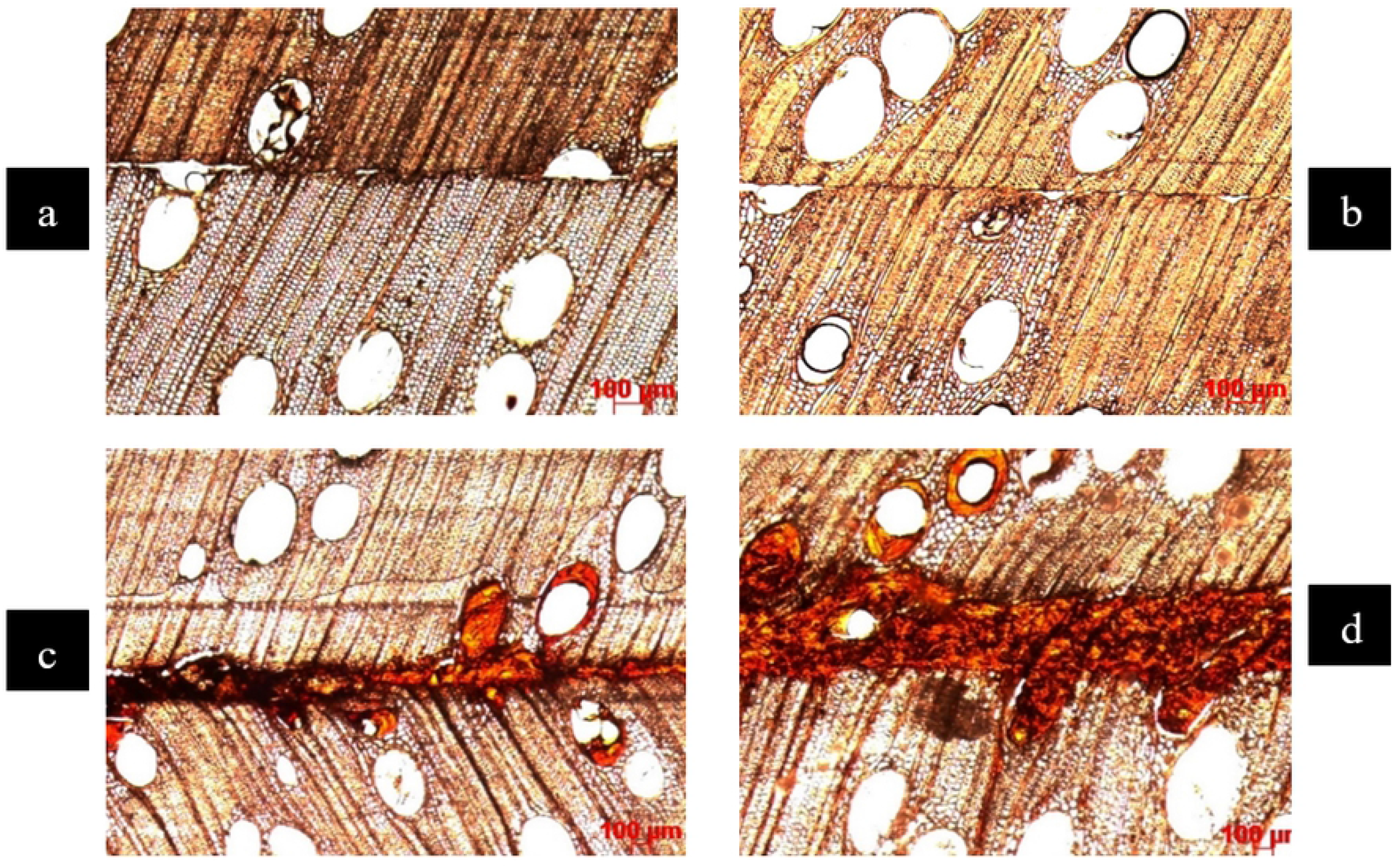
Glue line in the transverse surface of elements glued with castor polyurethane (CP) and resorcinol-formaldehyde (RF). a) CP in the wood lamina of Group 1 (< 0.58 g cm^−3^); b) CP in the timber lamina of Group 2 (≥ 0.58 g cm^−3^); c) RF in the timber lamina of Group 1 (<0.58 g cm^−3^); d) RF in the timber lamina of Group 2 (≥ 0.58 g cm^−3^).

After delamination test, glulam blocks of *Eucalyptus grandis* × *Eucalyptus urophylla* did not presented changes that compromised the integrity of wood-adhesive interface. Moisture, pressure and temperature variation were satisfactory to guarantee structural element integrity, as pointed out in the literature [17].

After visual analysis of glue lines conditions, specimens did not present delamination slits; which means 0% delamination.

### Relationship between main and secondary glue lines with apparent wood density

Based on F test (p ≤ 0.05), effects of apparent density, adhesives and their interaction were significant to thickness of the main glue line. Outcome interaction between apparent density and adhesives were spread out and analyzed (Table 2).

**Table 2.** Average numbers for thickness of main and secondary glue lines

| Density<br>(g cm<sup>-3</sup>) | Main glue line<br>(μm) |  | Secondary glue line<br>(μm) |  |
| --- | --- | --- | --- | --- |
|  | Adhesive |  |  |  |
|  | CP | RF | CP | RF |
| < 0.58 g cm<sup>-3</sup><br>(Group1) | 3.17Ab* | 125.22Ba | 51.77Ab | 278.18Ba |
| ≥ 0.58 g cm<sup>-3</sup><br>(Group 2) | 1.70Ab | 217.77Aa | 71.79Ab | 432.83Aa |

\* Horizontal and vertical lowercase letters do not differ from each other by variance analysis ( $F \leq 0.05$ ); CP: castor polyurethane adhesive; RF: resorcinol-formaldehyde adhesive.

When it comes to CP adhesive, the contrast between average apparent density levels was statistically zero. Therefore, glulam elements in Group 1 and 2 with CP adhesive showed no significant difference for both glue line thicknesses.

### Pearson’s correlation on studied variables

Main glue line thickness showed strong and significant correlation (t < 0.01) with adhesive (0.739) (Table 3). Correlation is a statistical analysis that measures the association between variables. Therefore, it was possible to say that the behavior presented by glue line thickness was expected due to the adhesive.

**Table 3.** Pearson correlation coefficients based on the average of studied variables for the hybrid *Eucalyptus grandis* x *Eucalyptus urophylla*.

|  | Denap# | Ades | LP | LS | DVaso | Hraio | Lraio | NCel |
| --- | --- | --- | --- | --- | --- | --- | --- | --- |
| **Denap** | 0 | 0 | 0.131** | 0.125** | 0.197** | 0.105<sup>ns</sup> | 0.235<sup>ns</sup> | 0.1394<sup>ns</sup> |
| **Ades** |  | 0 | 0.739** | 0.070<sup>ns</sup> | 0 | 0 | 0 | 0 |
| **LP** |  |  | 0 | 0.060<sup>ns</sup> | 0.100<sup>ns</sup> | -0.100<sup>ns</sup> | -0.050<sup>ns</sup> | -0.080<sup>ns</sup> |
| **LS** |  |  |  | 0 | -0.003<sup>ns</sup> | 0.023<sup>ns</sup> | 0.175** | -0.019<sup>ns</sup> |
| **DVaso** |  |  |  |  | 0 | 0.221** | -0.249<sup>ns</sup> | -0.104<sup>ns</sup> |
| **Hraio** |  |  |  |  |  | 0 | 0.361** | 0.503** |
| **Lraio** |  |  |  |  |  |  | 0 | 0.608** |
| **NCel** |  |  |  |  |  |  |  | 0 |

## Discussion

### Physical and anatomical characteristics for the hybrid *Eucalyptus grandis* × *Eucalyptus uroplylla*

The largest vessel diameters were found on Group 2 timber boards samples (ρ ≥ 0.58 g cm^−3^) (138.56 μm), value 7.57% higher than Group 1 (ρ < 0.58 g cm^−3^). Results found in this study were lower than reported in the literature [18, 19], respectively, 125.5 and 116 μm, for same hybrid trees.

In this study, used specie were 11 years old. Young to mature wood transition happens gradually between seven to fourteen years of growth for this specie [20]. Possibly, wood samples came from a more mature tree, with thick wood walls and reduced size of its cellular elements. This fact could explain the contrast between reduced values for vessels found in this work and those mentioned in literature.

Group 1 wood samples (lighter boards) presented 30.70% superior vascular frequency when compared to Group 2, respectively 7.82 and 7.76 vessels mm^−2^. The hybrid used in the research was provided in a batch, for this reason, it is possible that some timber boards, originally came from trees with different ages. Age is an important factor when it comes to adult wood formation, because it allows higher thickening of cell walls, vessels and parenchyma. Therefore, it may have influenced the measured difference in board anatomical elements [21]. Higher vascular frequency for Group 1 woods was compensated by smaller vessel diameter size. Woods with smaller vascular diameters have a higher vascular frequency [22].

Group 2 timber boards showed 7.80% higher rays height than boards from Group 1, respectively, 254.58 and 234.72 μm. Rays width in wood from Group 2 was 17.61% higher than Group 1 and the number of cells was 8.53% higher in Group 2. Different anatomical behaviors on *Eucalyptus* genus can be explained by silvicultural treatments, plant growth and tree age [22, 23].

### Adhesive-wood interface, delamination and glue line analysis

When conditions like wettability, roughness and cleanliness, or even surface treatment prior adhesive are ideal for gluing, adhesive penetration happens more deeply into wood capillary structure [24]. To be able to create a strong bonding between an adhesive and the substrate, it is required enough resin to penetrate into wood components, which means that the adhesive must have satisfactory mobility [24]. Thereby, glued elements possibly had ideal conditions in the surface to create a strong connection [7] and the amount of used adhesive was enough.

There was adhesive penetration into rays and vessels elements, which was possible due to perforations on their walls that allowed the inter-cells communication. These characteristics can help adhesive acquiring by the structure and in the anchoring formation between both [7].

Adhesive penetration into ray cells may have happened through transfer cells, since they are made of parenchymatic cells and they work as conducting elements in the radial direction. Parenchyma cells have a modified wall to allow transportation, which is generally short-distance [7]. The specie anatomical condition, high or low density, reflects on adhesive penetration [25, 26].

When adhesive has corrected viscosity and finds in substrate ideal conditions of surface and microscopic structure, the product created from this gluing must have equal or superior characteristics than the sum of individual characteristics from the materials that it was made from.

Adhesive penetration depth into wood may have helped in the resistance showed by the specimens in delamination test. Another factor that could explain this behavior are the inherent characteristics of used adhesives that have high resistance to humidity, which makes them able to be used outdoor [27, 33].

In this study, the resulting numbers for main and secondary glue line thickness were different from those found in species like *Eucalyptus cloeziana* according to literature [28]. This is associated with longer time and lower pressure applied in the present work, respectively 48 hours and 1 MPa.

RF adhesive presented opposite behavior to polyurethane adhesive in both glue lines formed, showing contrast between the average apparent density levels, statistically different from zero (F ≤ 0.05).

Highest thickness averages to main and secondary glue lines were found in Group 2, 42.50% and 35.73% higher than group 1 averages for RF (Table 2). These numbers are higher than those reported in previous studies for double glue lines 150 g cm^−2^ - 52.13 μm and 60.23 μm respectively [28].

Some variables are very important to the timber glueing process, among them, adhesive amount to be applied in each specie, wood blade thickness, pressure and pressing time [24, 29, 30].

There was a significant difference between glue lines glued with different adhesives. The highest averages were found in those elements glued with RF adhesive, in both groups. In Group 1, RF Glulam elements presented 97.47% averages (main glue line) and 81.39% (secondary glue line) higher than those glued with polyurethane castor adhesive. Group 2, the averages were also 99.22% and 83.41%, respectively, higher than polyurethane glued elements.

Anatomical characteristics are the reason why Group 2 showed higher numbers, because vessels with larger diameters, high and wide rays provided ideal conditions to adhesive mobility and penetration. However, the difference found between adhesives, can be explained by their distinct viscosity.

The higher the viscosity showed by the adhesive, the higher its difficulty of spreading, which is caused by the lower flowability, resulting in less adhesive penetration on the capillary structure of the wood [11].

### Mathematical correlation between physical and anatomical wood characteristics

When adhesive does not show ideal viscosity, there may be difficulties to get into porous structure of the wood, because of reduction or obstruction of empty spaces. This could lead to a thicker main glue line formation and lower depth penetration of the adhesive, which was not observed in this study.

Main glue line presented inverse and not significant correlation to ray characteristics, which means that these variables are not demanding when it comes to gluing. The necessary penetration of the adhesive to anchor in the macro and microscopic regions of the wood will be facilitated if the species shows characteristics such as higher porosity, large diameters vessels and high and wide rays, as observed in the thickness data of the glue line (Table 1).

A wood increase causes a reduction of the porosity, consequently, the wood will have a greater contact area and increase in the mechanical resistance to the worth which it is submitted.

The height of rays had a moderate correlation with cells number, also, rays width and number of cells, respectively 0.503 and 0.608, both under significant (t <0.01). Since the rays are formed by parenchyma cells and have an indeterminate length, it can be said that, increasing the height and width of rays will increase their capacity to create new storage cells [31], which can explain the correlation between these variables.

## Conclusions

– The anatomical structure showed by eucalyptus wood in this study, allowed the adhesive penetration into wood vessels and rays;
– The apparent density difference showed by the two groups gave distinction to the anatomical elements variability presented by timber boards;
– The adhesives used had satisfactory characteristics, what was proved by the delamination test;
– Tested adhesives are qualified for structural uses, since variations of humidity, pressure and temperature did not promoted weakness of the glue line;
– There was a significant interaction between glue line thickness and apparent density. Woods of higher apparent density (ρ ≥ 0.58 g cm^−3^) glued with RF adhesive responded better to this interaction;
– Rays height and width had a moderate correlation with cell numbers. This characteristic qualifies the wood as favorable to gluing, since larger rays will contribute in the adhesive permeability.

## Acknowledgement

This work was supported by “Foundation for Support Research and Innovation of Espírito Santo” (FAPES) and “National Council for Scientific and Technological Development” (CNPq).

